# Feminizing *Wolbachia* endosymbiont disrupts maternal sex chromosome inheritance in a butterfly species

**DOI:** 10.1101/115386

**Authors:** Daisuke Kageyama, Mizuki Ohno, Tatsushi Sasaki, Atsuo Yoshido, Tatsuro Konagaya, Akiya Jouraku, Seigo Kuwazaki, Hiroyuki Kanamori, Yuichi Katayose, Satoko Narita, Mai Miyata, Markus Riegler, Ken Sahara

**Affiliations:** Institute of Agrobiological Sciences, National Agriculture and Food Research Organization, 1-2, Owashi, Tsukuba, Ibaraki 305-0854, Japan; Laboratory of Applied Entomology, Faculty of Agriculture, Iwate University, 3-18-8, Ueda, Morioka 020-8550, Japan; Graduate School of Science, Kyoto University, Kitashirakawa Oiwake-cho, Sakyo-ku, Kyoto 606-8502, Japan; Institute of Crop Science, National Agriculture and Food Research Organization, 1-2, Owashi, Tsukuba, Ibaraki 305-0854, Japan; Tsukuba Primate Research Center, National Institute of Biomedical Innovation, Health and Nutrition, 1-1, Hachimandai, Tsukuba, Ibaraki 305-0843, Japan; Graduate School of Horticulture, Chiba University, Matsudo 648, Matsudo, Chiba 271-8510, Japan; Hawkesbury Institute for the Environment, Western Sydney University, Locked Bag 1797, Penrith, New South Wales 2751, Australia

**Author notes:** **For correspondence:** (DK); (KS).

## Abstract

Genomes are vulnerable to selfish genetic elements that enhance their own transmission often at the expense of host fitness. Examples are cytoplasmic elements such as maternally inherited bacteria that cause feminization, male-killing, parthenogenesis and cytoplasmic incompatibility. We demonstrate, for the first time, that segregation distortion, a phenomenon so far seen only for nuclear genetic elements, can also be caused by a cytoplasmic element, the ubiquitous endosymbiotic bacterium *Wolbachia*. For *Eurema mandarina* butterfly lineages with a Z0 sex chromosome constitution, we provide direct and conclusive evidence that *Wolbachia* induces production of all-female progeny by a dual role: the compensation for the female-determining function that is absent in Z0 lineages (feminization) and the prevention of maternal sex chromosome inheritance to offspring as a specific type of segregation distortion. Therefore, our findings highlight that both sex determination and chromosome inheritance — crucially important developmental processes of higher eukaryotes — can be manipulated by cytoplasmic parasites.

## Introduction

Genomes of sexually reproducing organisms are exposed to genetic conflicts. For example, some genes bias reproduction towards male offspring while other genes within the same genome may favor reproduction of more daughters. Selfish genetic elements (SGEs), such as meiotic drivers, cytoplasmic sex ratio distorters and transposons, are extreme examples, which enhance their own transmission often at the expense of their hosts’ fitness [1,2]. There is growing evidence that SGEs, and their genetic conflict with host genomes, trigger important evolutionary change and innovation in eukaryotes [2].

Segregation distortion (SD), also referred to as meiotic drive, is a violation of Mendelian law as it leads to the more frequent inheritance of one copy of a gene than the expected 50% [3,4]. A segregation distorter that sits on a sex chromosome biases the sex ratio. For example, X-linked segregation distorter (X drive) and Y-linked segregation distorter (Y drive) in flies (Diptera), result in female-biased and male-biased sex ratios, respectively [4]. In male-heterogametic species, X and Y segregation distorters are expected to be encoded in the nuclear genome. In female-heterogametic species, however, W chromosome and cytoplasm behave as a single linkage group and thus distortion of sex chromosome inheritance in female-heterogametic species can theoretically also be caused by cytoplasmic elements. Although this possibility has previously been proposed [5,6], lack of empirical evidence questions whether it is mechanistically possible for cytoplasmic elements to cause SD.

*Wolbachia pipientis* (Alphaproteobacteria), simply referred to as *Wolbachia*, attracts significant interest in evolutionary and developmental biology but also in applied fields such as pest management because it can manipulate reproduction of arthropods in various ways such as cytoplasmic incompatibility, parthenogenesis induction, feminization and male-killing [7]. Here we demonstrate for the first time that *Wolbachia* is responsible for the disruption of sex chromosome inheritance, which can also be seen as a form of segregation distortion, in any host species. We do this by providing multifaceted and conclusive evidence that in the butterfly *Eurema mandarina Wolbachia*-induced SD is the underlying mechanism for the production of all-female progeny. In most populations, *E. mandarina* is infected with the cytoplasmic-incompatibility (CI)-inducing *Wolbachia* strain *w*CI at a high prevalence of close to 100% [8,9]. Hiroki et al. [10,11] first reported all-female offspring production in *E. mandarina* (then known as *Eurema hecabe* yellow type), which was considered to be due to the feminization of genetic males (ZZ) by co-infections with the *Wolbachia* strain *w*Fem (hereafter referred to as double infection CF while single infection with *w*CI is referred to as C). Three observations about CF lineages supported this view, i.e., (a) antibiotic treatment of adult females led to the production of all-male offspring [10], (b) antibiotic treatment of larvae resulted in intersex adults [12] and (c) females did not have the W chromatin body [10,12]. This has recently been challenged, because it was demonstrated that CF females have only one Z chromosome and that this Z chromosome always derived from their fathers implying that a SD mechanism may be in place albeit it was not clear whether *Wolbachia* induced this SD [13]. As a consequence two novel (yet untested) hypotheses were formed, namely, that CF females have either a Z0 or a W’Z sex chromosome set (whereby W’ cannot be visualized in W chromatin assays and does not have a female-determining function), and that the disruption of Z chromosome inheritance occurs in CF lineages due to *Wolbachia* or another factor, such as those encoded by the host nucleus.

In a multifaceted approach, by combining fluorescence in situ hybridization (FISH), genome sequencing, quantitative PCR, reverse transcription PCR and antibiotic treatment, we have tested these two hypotheses and revealed that CF females are Z0, and that *Wolbachia* is the cause for both the disruption of Z chromosome inheritance and the feminization of Z0 individuals. Our results demonstrate, for the first time, *Wolbachia* as the agent that is responsible for distorted sex chromosome inheritance, and thereby highlight that cytoplasmic elements can have profound effects on oogenesis, sex chromosome inheritance and sex determination – fundamental biological processes of eukaryotes.

## Results

### All-female-producing CF females have a Z0 sex chromosome constitution

We performed FISH on *E. mandarina* chromosomes prepared from CF females, C females, and C males collected on Tanegashima Island (***Figure 1***; ***Figure 1—figure supplement 1***). In the mitotic complement of C females, which harbor a 2*n* = 62 karyotype, genomic probes highlighted the W chromosome, with scattered signals on the other chromosomes (***Figure 2A***; see Materials and Methods for technical details). A probe for the Z-linked gene *Kettin* (*Ket*) identified the single Z chromosome in C females (***Figure 2A***), and also hybridized to the Z chromosome paired with the W chromosome in pachytene bivalents (***Figure 2J***). The *Ket* probe identified two Z chromosomes in the mitotic complement of C males (***Figure 2B***; 2*n* = 62). No painted W chromosome was observed in interphase nuclei (***Figure 2H, I***), the mitotic complement (***Figure 2C***) and pachytene complement (***Figure 2L***) of CF females, but the *Ket* signal appeared on the single Z chromosome in the mitotic complement (***Figure 2C***) and Z univalent in the pachytene complement (***Figure 2L***). Based on the relative read counts homologous to *Bombyx mori* Z-linked and autosomal genes in females and males, our genome sequencing data support the notion that CF and C females have one Z chromosome (***Figures 2M–O***; ***Figure 2—figure supplement 1***), which is consistent with genomic qPCR data based on two loci, *Triosephosphate isomerase* (*Tpi*) and *Ket*, relative to the autosomal gene *EF-1α*[13]. Thus, our results directly reveal the sex chromosome constitution of C females, C males, and CF females as WZ, ZZ, and Z0, respectively. This confirms one of two previously suggested sex chromosome constitution of CF females [13] while it disproves another previous interpretation based on W-body diagnosis that CF females are ZZ [10,12].

**Figure 1.**
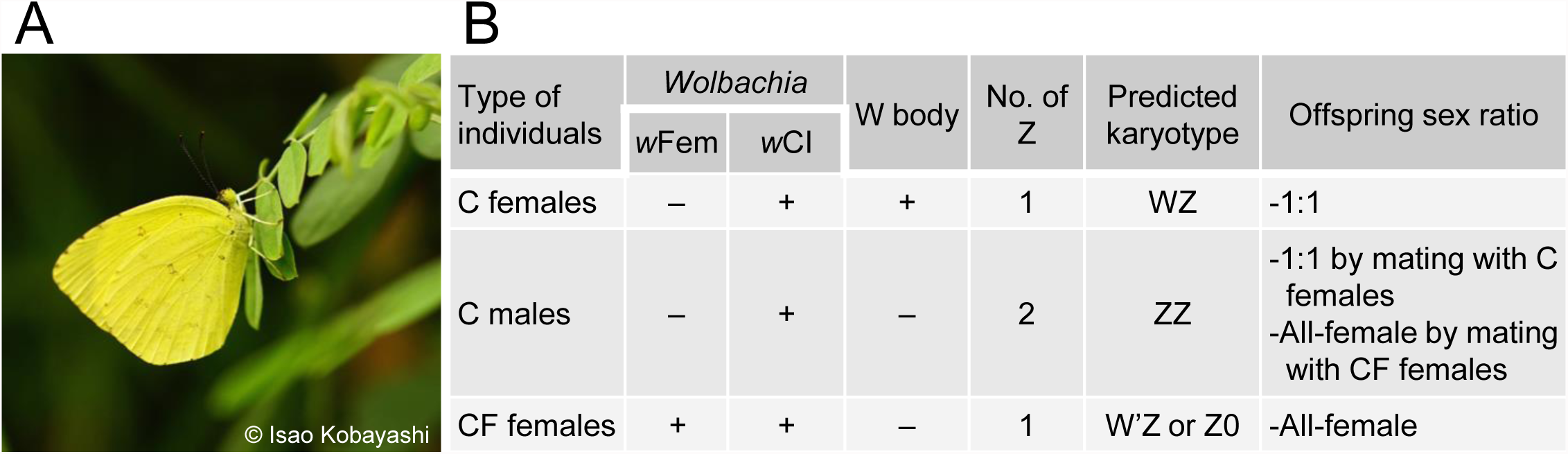
*E. mandarina* butterflies used in this study. (**A**) A photo of *E. mandarina* taken in Tanegashima Island. (**B**) Characteristics of three types of *E. mandarina* individuals inhabiting Tanegashima Island.

**Figure 2.**
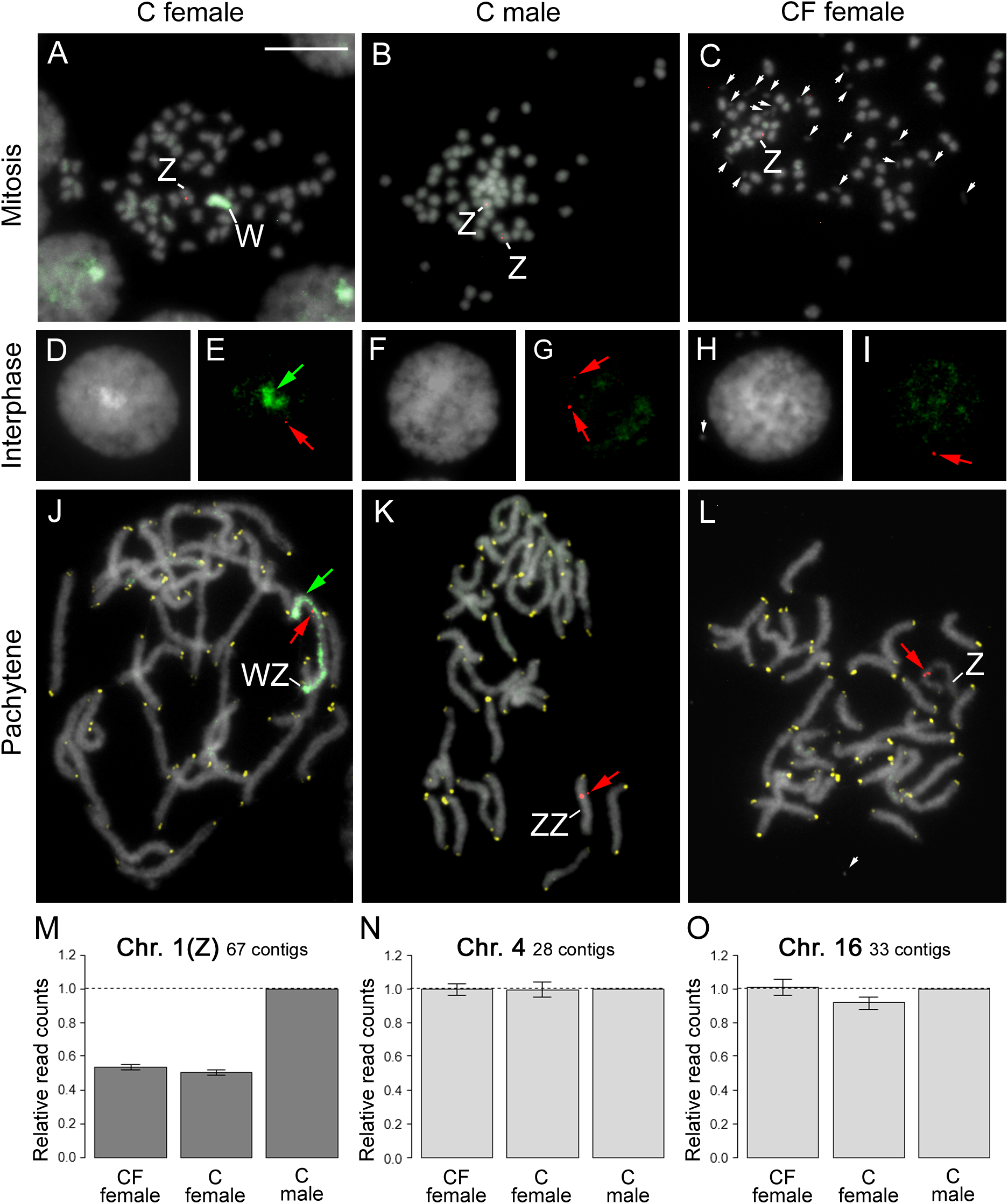
Fluorescence *in-situ* hybridization and sequence read counts for a C female, C male, and CF female *E. mandarina*. **A**–**C**: Mitotic complements hybridized with a genomic probe (green; green arrows) and a Z-linked *Ket* probe (red; red arrows) in a C female (2n = 62) (**A**), C male (2n = 62) (**B**), and CF female (2n = 61) (**C**). **D**–**I**: Genomic *in situ* hybridization (GISH) and FISH with a Z-linked *Ket* probe performed on interphase nuclei of *E. mandarina* C females (**D**, **E**), C males (**F**, **G**), and CF females (**H**, **I**). **J**–**L**: GISH, telomere-FISH and FISH with *Ket* probe performed on pachytene complements of *E. mandarina* C females (**G**, n = 31), C males (**H**, n = 31), and CF females (**I**, n = 31). Green paint signals in **A**, **E** and **J** revealed that C females have the W chromosome. The *Ket* probe signals (red) appeared on the Z pairing to the W in C females (**J**), the ZZ bivalent in C males (**K**), and the Z univalent of CF females (**L**). The single signals were observed both in C and CF female nuclei. The signals in C females (**J**) and males (**K**) clearly showed their respective WZ and ZZ chromosome sets, and a Z0 chromosome set in CF females (**L**). W: W chromosome; Z: Z chromosome; white arrows: *Wolbachia*-like structures. A bar represents 10 m. **M**–**O**: Relative normalized sequence read counts in CF females, C females, and C males for 67 contigs homologous to *Bombyx mori* loci on chromosome 1 (Z chromosome; **M**), 28 contigs homologous to *B. mori* loci on chromosome 4 (**N**), and 33 contigs homologous to *B. mori* loci on chromosome 16 (**O**), with relative read counts set to 1 (males). Details about genome sequencing are provided in Materials and Methods.

### All embryos oviposited by CF females are Z0

We performed real-time genomic qPCR (to detect Z-linked *Tpi* or *Ket* relative to autosomal *EF-1α*) on individual fertilized eggs, and found that C females oviposited embryos with either one or two Z chromosomes at nearly equal frequencies (***Figure 3A left***; ***Figure 3—figure supplement 1***). In contrast, all embryos oviposited by CF females were single Z carriers (***Figure 3A middle***; ***Figure 3—figure supplement 1***). These findings indicate that the progeny of CF females are exclusively Z0 individuals, supporting the view that the maternal Z chromosomes are not inherited in CF lineages.

**Figure 3.**
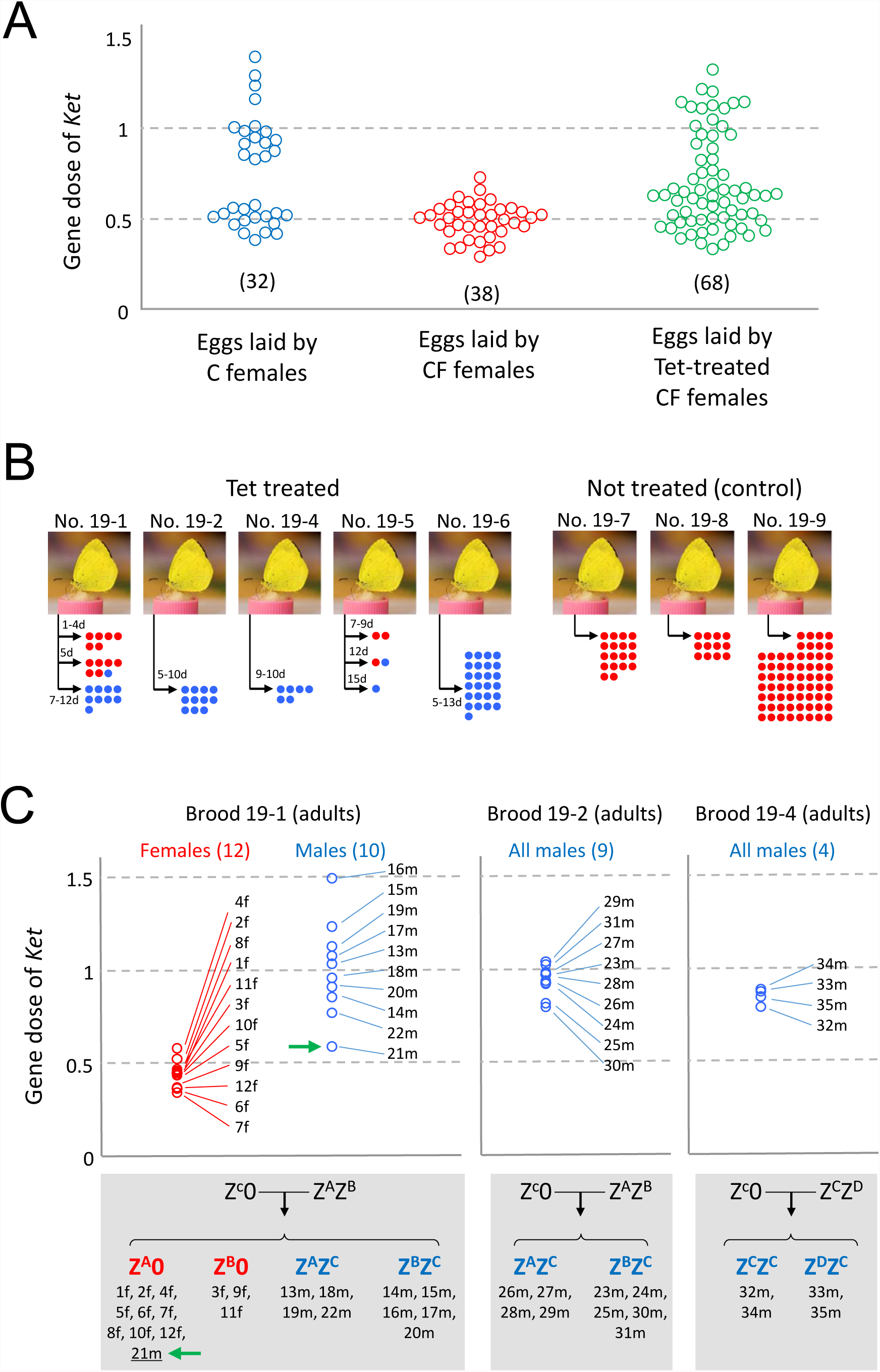
Effects of *w*Fem on Z-linked gene dose in *E. mandarina* offspring. (**A**) Estimate of the gene dose of *Ket* (relative gene copies per copy of *EF-1α*) by genomic quantitative polymerase chain reaction (qPCR) analysis in each of the fertilized eggs laid by C females, CF females, and tetracycline (tet)-treated CF females. Each colored circle represents a single fertilized egg. Sample sizes are given in parentheses. (**B**) Offspring sex ratio of five females tet-treated prior to oviposition and three non-treated CF females. Numbers to the left of the arrows represent duration (days) of tet treatment. Blue dots and red dots represent males and females, respectively. (**C**) Estimate of the gene dose of *Ket* (relative gene copies per copy of *EF-1α*) by genomic qPCR in each of the adult offspring produced by CF females that were tet-treated during the adult stage (prior to oviposition). Each circle represents an adult offspring. Z chromosomes of these offspring individuals were genotyped as Z^A^, Z^B^, Z^C^ or Z^D^ on the basis of intron nucleotide sequence of Z-linked *Tpi*. The green arrow points to a male individual (adult) whose karyotype was considered to be Z0 but possibly ZZ’ (see text for details). f: female, m: male.

### *Wolbachia* causes the exclusive production of Z0 embryos by CF females

To abolish the effects of *Wolbachia*, tetracycline (tet) was administered to adult CF females previously inseminated by antibiotic-treated male offspring of C females. The Z-linked gene dose of embryos laid by these tet-treated females ranged from approximately 0.5–1.0, indicating that some embryos are Z0 and others are ZZ (***Figure 3A right***; ***Figure 3—figure supplement 1***). This suggests that the *Wolbachia* strain *w*Fem in CF females causes the exclusive production of gametes without sex chromosomes that then develop as Z0 embryos after fertilization. Therefore, our finding is the first empirical evidence that in a female-heterogametic species the sex-specific linkage disequilibrium can be caused by cytoplasmic elements [5,6]. Furthermore, *Wolbachia*-like structures were observed near the chromosomes in CF females while less apparent in C females and C males, and this may represent different tropism and function of *w*Fem when contrasted with *w*CI (***Figure 2C***). Sixty-nine adults (15 females and 54 males) were obtained from offspring produced by five tet-treated adult CF females (***Figure 3B***). Three of these tet-treated females produced only male offspring. Exclusive production of males was previously observed in tet-treated *E. mandarina* females derived from a different population on Okinawa-jima Island, Okinawa Prefecture, Japan [10]. In this study, we obtained 15 female offspring from two broods in the first days after tet treatment; however, the mothers produced more males as the duration of tet treatment increased, and eventually produced only males. Examination of the Z-linked gene dose of these offspring by genomic qPCR showed that the females had one Z chromosome, whereas almost all of the males had two Z chromosomes (***Figure 3C***). The nucleotide sequences of the introns of the *Tpi* gene demonstrate that, in brood 19-1, all females (*n* = 12) were hemizygous and nine out of 10 males were heterozygous (***Figure 3C***; ***Figure 3—figure supplement 2***). Curiously, one male (21m) that exhibited the lowest gene dose of *Ket* (0.588) appeared to be hemizygous (***Figure 3C***). These results suggest that the emerged females had a Z0 sex chromosome constitution, whereas most males had a ZZ sex chromosome constitution, with one exception (21m) of either Z0 or ZZ’ (Z’ represents partial deletion/mutation in Z). These results also demonstrate that, in principle, tet-treated adult CF females can oviposit embryos with either a Z0 or ZZ sex chromosome constitutions (***Figure 3A right***). However, Z0 individuals appear to have zero or very low survival rates because few emerge as adults.

### Involvement of *Wolbachia* in the sex determination of *Eurema mandarina*

Next, we fed CF larvae a tet-containing diet. As previously observed [12], all individuals treated in this way developed an intersex phenotype at the adult stage, typically represented with male-like wing color and an incomplete male-specific structure on the wing surface (***Figure 4E and H***; ***Figure 4—figure supplement 2***). The qPCR assay to assess the Z-linked gene dose revealed that these intersexes (*n* = 23) had just one Z chromosome (***Figure 4I***), and therefore a Z0 genotype. Because these Z0 individuals were destined to develop as females without tet treatment, *w*Fem is likely to be responsible for female sex determination. Further evidence in support of this idea was obtained by examining the sex-specific splicing products of *dsx* (***Figure 4—figure supplement 3***), a widely conserved gene responsible for sexual differentiation [14]. Similar to *B. mori* [15], C females exhibited female-specific splicing products of *E. mandarina dsx* (*Emdsx^F^*), whereas C males had a male-specific splicing product of *E. mandarina dsx* (*Emdsx^M^*; Lanes 1 and 2 in ***Figure 4A***, respectively; ***Figure 4B***). Similarly to C females, CF females exhibited exclusive expression of *Emdsx^F^* (Lanes 3 and 4 in ***Figure 4A***; ***Figure 4B***). Intersexual butterflies, generated by feeding the larval offspring of CF females a tet-containing larval diet, expressed both *Emdsx^F^* and *Emdsx^M^* (Lanes 5 and 6 in ***Figure 4A***; ***Figure 4—figure supplement 1***).

**Figure 4.**
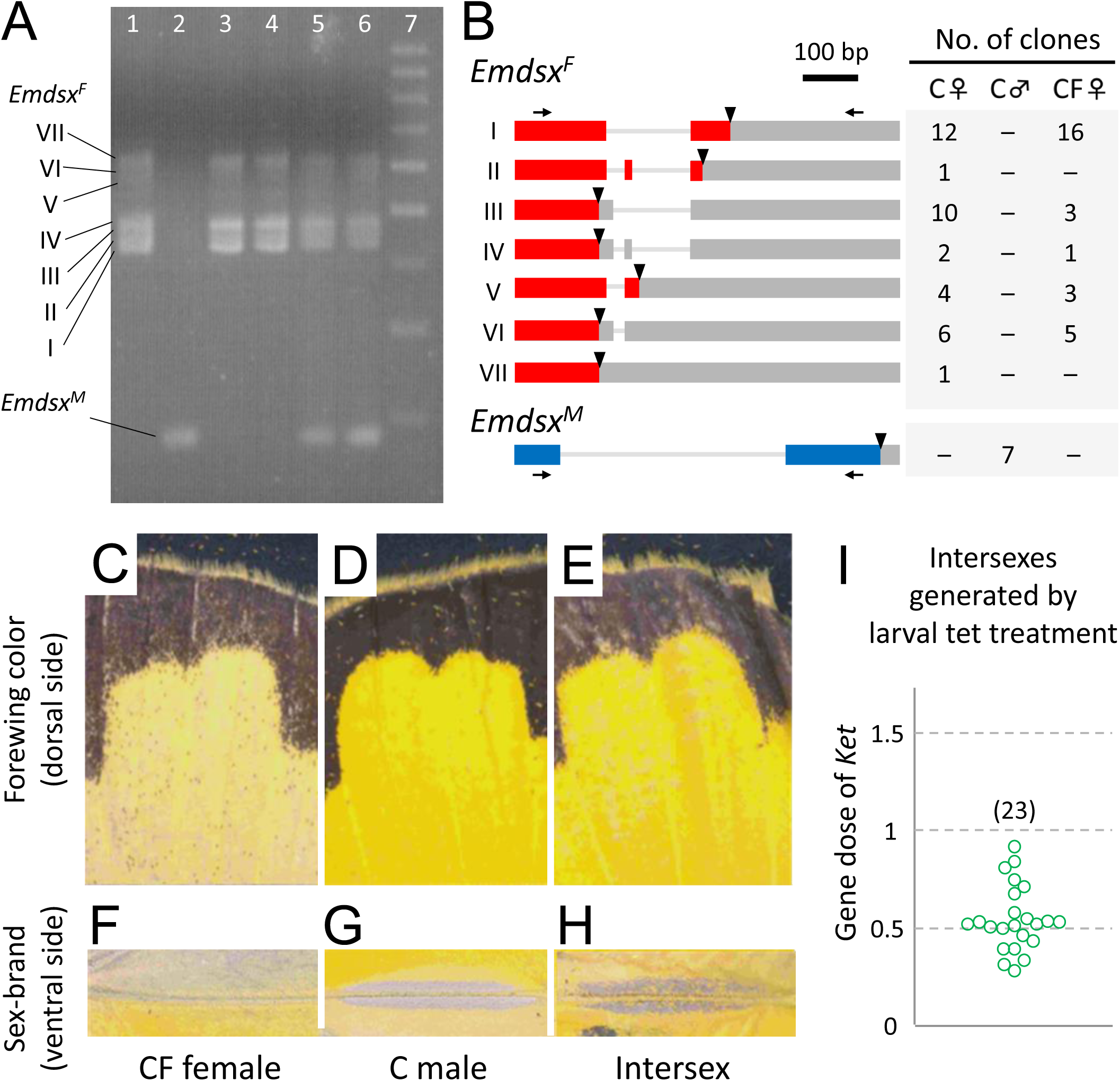
Effects of *w*Fem on splicing of the *doublesex* gene in *E. mandarina*.(**A**) Reverse-transcription polymerase chain reaction (RT-PCR) products of *E. mandarina doublesex* (*Emdsx*) run on an agarose gel. Lane 1: C female; lane 2: C male; lanes 3 and 4: CF females; lanes 5 and 6: intersexes generated by tetracycline (tet) treatment of larvae produced by CF females; lane 7: 100-bp ladder. Females have at least seven splicing products, whereas males have a single product. (**B**) Structures of the splicing products of *Emdsx*. Translated regions are indicated by red and blue bars, untranslated regions by gray bars, and stop codons by triangles. Numbers of clones obtained by cloning the RT-PCR products are shown in the table on the right. **C**–**H**: color and morphology of forewings. Females are pale yellow on the dorsal side of the forewings (**C**) and do not have sex brand on the ventral side of the forewings (**F**), while males are intense yellow on the dorsal side of the forewings (**D**) and have sex brand on the ventral side of the forewings (**G**). Many of the intersexes generated by tet-treating CF larvae are strong yellow on the dorsal side of the forewings (**E**) and have faint sex brand on the ventral side of the forewings (**H**).

## Discussion

We provide comprehensive and conclusive indirect (qPCR of Z gene dosage) and direct (W chromosome painting; genomic analyses) evidence for the loss of the W chromosome from CF individuals. Furthermore, we demonstrate that the *Wolbachia* strain *w*Fem is directly responsible for chromosomal segregation distortion (SD) by causing the disruption of sex chromosome inheritance in CF females of *E. mandarina*. This is the first empirical proof for previous theoretical predictions that cytoplasmic SGEs, such as *Wolbachia*, can cause SD. In *E. mandarina*, *w*Fem has a dual role in both causing segregation distortion and feminization in Z0 lineages that have lost W chromosome and its feminizing function.

### *Wolbachia* disrupts Z chromosome inheritance in Z0 females

Our data provides evidence that the exclusive production of Z0 embryos by CF females is due to a yet unidentified developmental process that leads to the disruption of sex chromosome inheritance in CF females prior to oviposition, thereby the absence of maternal Z chromosome in CF offspring. This process can be referred to as SD, according to established conceptual frameworks [5,6]. We believe that two mutually exclusive hypotheses can account for the SD observed in CF individuals (***Figure 5A***). The first assumes that a gamete without the maternal Z chromosome (or without any sex chromosome overall), is always selected to become an egg pronucleus (meiotic drive *sensu stricto*) (***Figure 5A left***) [16]. The second assumes that meiosis itself is normal, and that maternal Z chromosomes (or sex chromosomes in general), are selectively eliminated from Z-bearing gametes during, or possibly after, meiosis (***Figure 5A right***). At present, it is unclear which of the two scenarios (meiotic drive *sensu stricto* or elimination of the maternal Z at a later stage) is more plausible. However, it is noteworthy that, in the moth *Abraxas grossulariata*, a matriline consisting of putative Z0 females was observed to produce only females or a great excess of females, and the underlying mechanism was considered to be the selective elimination of Z chromosomes [17–20]. However, the presence of cytoplasmic bacteria such as *Wolbachia* has not yet been examined for this moth species. If we assume that the elimination of the maternal Z chromosome is the mechanism of the SD in *E. mandarina*, the exceptional individual 21m (***Figure 3C***) could be viewed as ZZ’ rather than Z0, wherein Z’ is a maternal Z chromosome that was only partially deleted in the position including *Tpi* and *Ket* by the incomplete action of *w*Fem. It is possible to further speculate that the presence of *w*Fem results in the elimination of sex chromosomes in general (Z or W chromosomes) and, therefore, the absence of W chromosomes in CF females may also be a direct effect of *w*Fem.

**Figure 5.**
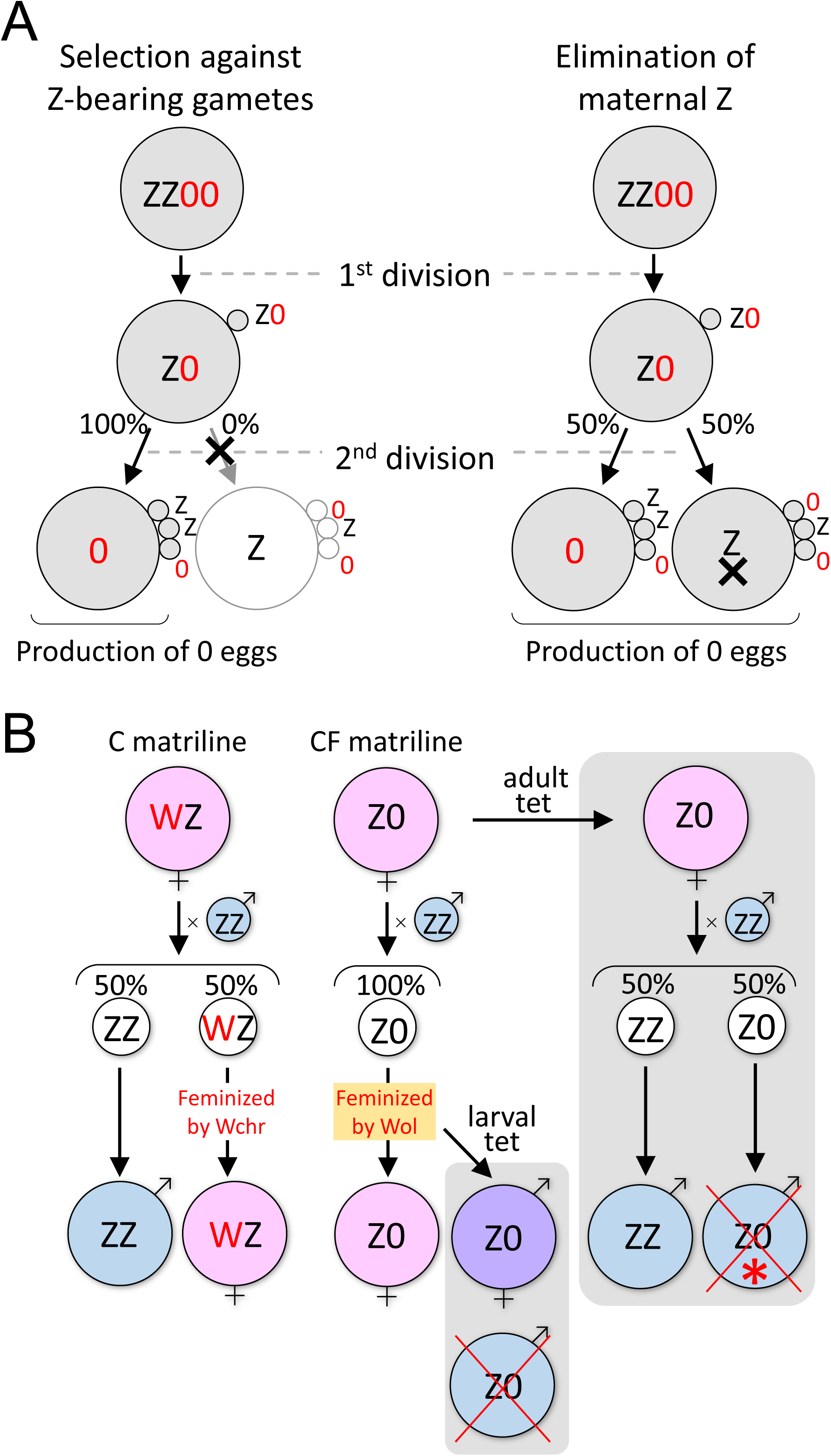
(**A**) Schematic illustration of two alternative mechanistic models of sex-chromosome segregation distortion that explain the observed data. The “Selection against Z gametes” model assumes that Z-bearing gametes are selected against during meiosis (left). The “Elimination of maternal Z” model assumes that Z chromosomes are eliminated during or after normal meiosis, while all the autosomes being intact (right). (**B**) All-female production explained by *Wolbachia*–host interaction. Effects of *w*Fem on the development and sex determination of *E. mandarina*, and outcomes of larval versus adult tet treatment are illustrated. Asterisk: The majority of Z0 males die, but a few survived.

### The feminizing effect of *Wolbachia* compensates for the loss of the W chromosome in Z0 individuals

In general, lepidopterans species with Z0/ZZ sex chromosome constitution are considered to determine their sexes by Z-counting mechanisms, wherein ZZ is male and Z0 is female [21,22]. However, the appearance of the male phenotype in Z0 individuals of *E. mandarina* after antibiotic treatment suggests that *w*Fem in Z0 individuals compensates for the loss of W and its feminizing function (***Figure 5B***). We speculate that the W chromosome of *E. mandarina* acts as an epistatic feminizer. In *B. mori*, the W chromosome – more specifically, a piRNA located on the W chromosome – acts as an epistatic feminizer by silencing *Masculinizer* on the Z chromosome [23].

Reduced survival of Z0 individuals or their offspring after antibiotic treatment of larvae or adults, respectively, may suggest improper dosage compensation in Z0 males. Improper dosage compensation was also proposed to be the cause of male- and female-specific lethality in *Wolbachia*-infected and cured lines of *Ostrinia* moths [24–27].

### How did the coordinated dual effects of *Wolbachia* evolve?

We demonstrated that *w*Fem causes SD and feminization in *E. mandarina* in two steps (***Figure 5B***). This is similar to the dual role of *Wolbachia* and *Cardinium* in haplodiploid parasitoid wasps where they induce thelytokous parthenogenesis in a two-step mechanism, comprising diploidization of the unfertilized egg followed by feminization [28,29]. Here, we develop the potential evolutionary scenario that led to the appearance of both effects in *E. mandarina* (***Figure 6***). A WZ female *Eurema* butterfly may have acquired *w*Fem that exerted a feminizing effect on ZZ males. The feminizing effect was lethal to ZZ individuals because of improper dosage compensation, as evident in *Wolbachia*-infected *Ostrinia* moths (***Figure 6A***) [26,27]. This could be viewed as a manipulation similar to a male-killing phenotype [30,31]. However, the feminizing effect of *w*Fem was redundant in WZ females where the W chromosome acted as a female determiner [23]. Conversely, the function of W had also become redundant in CF individuals and this could have led to the loss of the W chromosome and the rise of a Z0 lineage (***Figure 6B***). Similarly, in *Ostrinia* moths, a female-determining function is thought to have been lost from the W chromosome in *Wolbachia*-infected matrilines [25]. Spontaneous loss of a nonfunctional W chromosome may be easier than expected: in a wild silkmoth *Samia cynthia*, the W chromosome does not have a sex-determining function, and Z0 females are frequently obtained in experimental crosses between subspecies [32]. *Wolbachia* has previously been found to be involved in the loss and birth of W chromosomes in the woodlouse *Armadillidium vulgare* [33,34]. However, in *A. vulgare* it has not yet been tested whether *Wolbachia* interferes with chromosome segregation and inheritance as we have mechanistically demonstrated it for *E. mandarina*; i.e., after the loss of the W chromosome in CF lineages, *Wolbachia* then acquired a novel function that affected female oogenesis and resulted in SD (***Figure 6C***). It is unlikely that SD arose prior to the feminization function of *Wolbachia*: if the appearance of SD were to precede the loss of the W chromosome, the feminizing or female-determining function would become unnecessary for *Wolbachia* because there would be no males. In the short term, disruption of Z chromosome inheritance in females in a female-heterogametic species represents a great advantage to cytoplasmic symbionts because all vertically transmitted symbionts gain the opportunity to survive. However, males are still required for fertilization, and fixation of the symbionts in the host population will inevitably lead to the extinction of both the symbionts and the hosts [35]. In the long term, suppressors against sex ratio distortion, as has been observed for the male-killing phenotypes in the butterfly *Hypolimnas bolina* or a ladybird beetle [36,37], can be expected to evolve in the host. However, the evolutionary outcomes of the suppression of a combined SD and feminization would be different from that of male-killing suppression, because it would lead to all-male progeny, resulting in the loss of the matriline that inherits the feminizing and sex-distorting *Wolbachia*. This process thereby selects for an increased frequency of WZ females.

**Figure 6.**
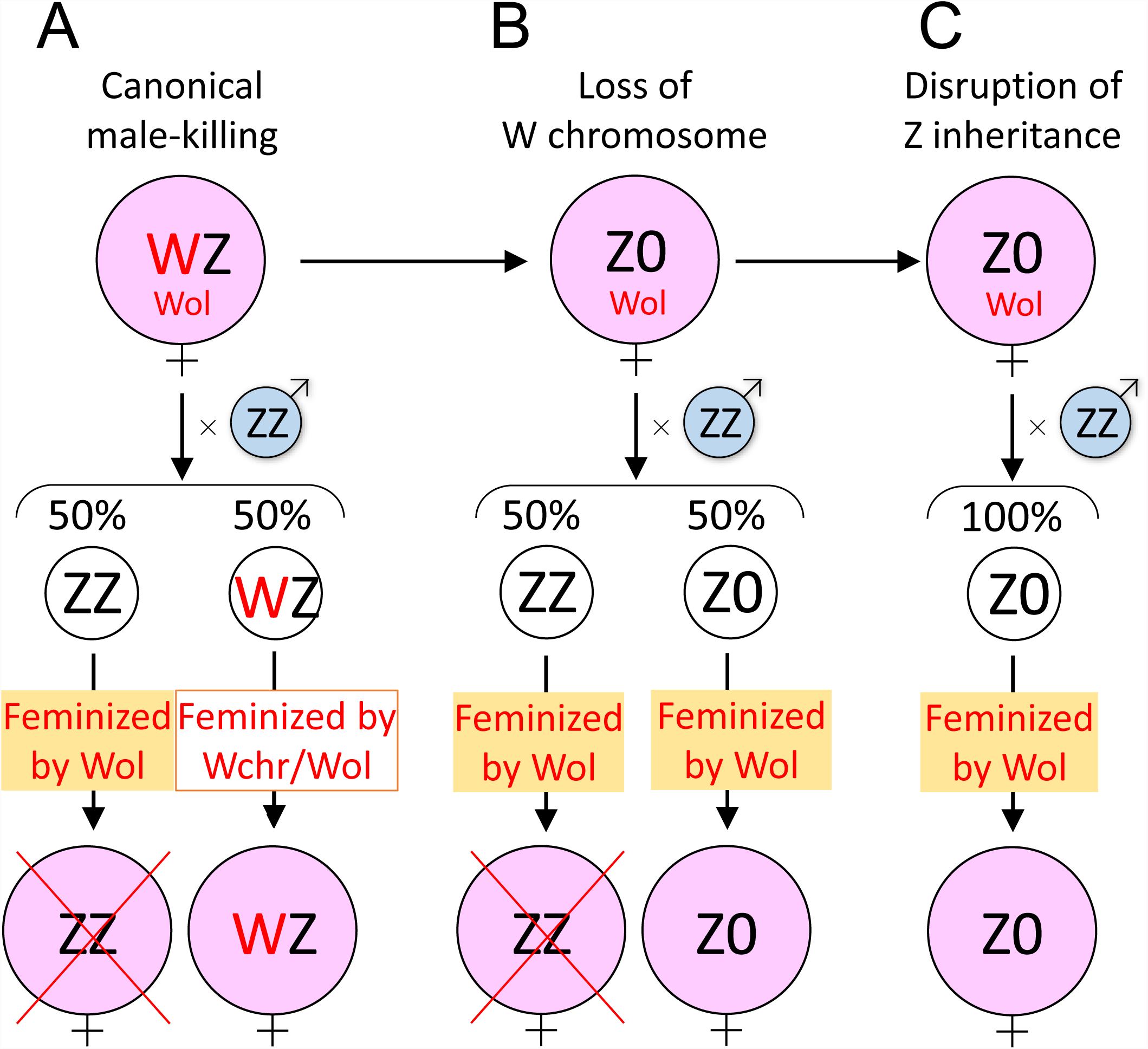
Hypothetical evolutionary trajectory of the *Wolbachia*–host interaction in *E. mandarina*. See Discussion for details.

## Concluding remarks

In summary, we demonstrate for the first time that the manipulation of sex chromosome inheritance and cytoplasmically induced SD can be added to the repertoire of host manipulations induced by *Wolbachia*. Therefore, the host effects of this bacterium are far more diverse and profound than previously appreciated. Disentangling these complex interactions between insects and *Wolbachia* may provide further exciting discoveries in the areas of host–parasite interactions, endosymbiosis as well as cell and chromosome biology in years to come, and perhaps also provide new avenues for pest population control.

## Materials and methods

### Collection and rearing of *E. mandarina*

Female adults of *E. mandarina* (Lepidoptera: Pieridae) were collected on Tanegashima Island, Kagoshima, Japan (***Figure 1—figure supplement 1***). In the laboratory, each female was allowed to lay embryos on fresh leaves of *Lespedeza cuneata* (Fabales: Fabaceae) in a plastic cup with absorbent cotton immersed with 5% honey solution. The artificial diet for larvae was prepared by mixing leaf powder of *Albizia julibrissin* (Fabales: Fabaceae) in the custom-made Silkmate (Nihon-Nosa, Yokohama, Japan) devoid of mulberry leaves. Insects were reared under the 16 h/8 h light /dark photoperiod at 25°C.

### Antibiotic treatment

We performed antibiotic treatment of two different stages (larval stage and adult stage) of *E. mandarina*. For larval antibiotic treatment, larvae were fed with the artificial diet (shown above) containing 0.05% tetracycline hydrochloride (tet). For adult antibiotic treatment, female adults were fed with 5% honey solution containing 0.1% tet. Specifically, CF females were mated to antibiotic-treated male offspring of C females. Antibiotic treatment of these males was performed in the larval stage and prevented CI in the crossing. After mating, each CF female was allowed to lay embryos on fresh leaves of *L. cuneata* in a plastic cup with absorbent cotton immersed with 5% honey solution containing 0.1% tet. Fresh leaves of *L. cuneata* and cotton with tet-containing honey solution were exchanged daily.

### Diagnosis of *Wobachia* strains

To diagnose *Wolbachia* strains in *E. mandarina*, several legs of each adult were homogenized in STE buffer (10 mM Tris-HCl (pH 8.0), 1 mM EDTA (pH 8.0), 150 mM NaCl) and incubated at 56°C for 30 min followed by 92°C for 5 min. After centrifugation at 15,000 rpm for 2 min, the supernatant was used for polymerase chain reaction (PCR) using different primer pairs. The primer pair wsp81F (5′–TGGTCCAATAAGTGATGAAGAAAC–3′) and wsp691R (5′–AAAAATTAAACGCTACTCCA–3′) amplifies a ca. 610-bp fragment of the *Wolbachia wsp* gene [38]. The primer pair wsp81F and HecCIR (5′–ACTAACGTCGTTTTTGTTTAG–3′) amplifies a 232-bp fragment of the *wsp* gene of *w*CI, while the primer pair HecFemF (5′–TTACTCACAATTGGCTAAAGAT–3′) and the wsp691R amplifies a 398-bp fragment of *wsp* gene of *w*Fem [11,39].

### Whole genome sequencing and de novo assembly

We performed whole genome sequencing for three types of *E. mandarina* individuals (CF females, C females and C males) that were collected on Tanegashima Island, Japan (***Figure 1—figure supplement 1***). Six genomic DNA libraries (two libraries for each sample type derived from two individuals) were constructed following manufacturer’s instructions (http://www.illumina.com). The average insert size of the libraries was approximately 350 bp and each library was multiplexed using a single indexing protocol. The genomic DNA libraries were sequenced by Illumina MiSeq using MiSeq Reagent Kit v3 (600-cycle) (Illumina, San Diego, CA). Generated raw reads (8.31 Gb, 5.34 Gb, and 6.94 Gb for CF females, C females and C males, respectively) were filtered by Trimmomatic [40] and then mapped to the complete genome of *Wolbachia* strain *w*Pip (GenBank: NC_010981.1) by Bowtie2 [41]. Mapped reads were discarded and then remaining reads of the three samples were merged and de novo assembled by SGA assembler [42]. Generated genome contig sequences were used for further analysis.

### Analysis of mapped read counts on chromosomes

To verify that CF and C females have one Z chromosome, we compared normalized mapped read counts of the three samples on Z chromosomes and remaining chromosomes. The filtered reads of each sample were mapped to the genome contigs by Bowtie2 (only concordantly and uniquely mapped reads were counted) and then normalized mapped read count of each sample on each contig was calculated based on the ratio of the number of total mapped reads between the three samples. Nucleotide sequences of relatively long genome contigs (length is 2 kb or more) with enough coverage (20 or more mapped reads) were extracted and compared with the gene set A of *B. mori* [43] by blastx search (cutoff e-value is 1e-50). Genome contigs with blastx hits were extracted and classified into 28 chromosomes based on the location of the homologous *B. mori* genes. For each chromosome, the average number of relative normalized mapped read counts was calculated for each sample (the number of C males was normalized to 1) using the normalized mapped read counts in the classified genome contigs, respectively.

### Sanger sequencing

To genotype Z chromosomes, a highly variable intron of Z-linked triosephosphate isomerase (*Tpi*) gene was PCR amplified using the primers, 5′–GGTCACTCTGAAAGGAGAACCACTTT–3’ and 5′–CACAACATTTGCCCAGTTGTTGCAA–3′, located in coding regions [44]. The PCR products were treated with ExoSAP-IT^®^ (Affymetrix Inc., Santa Clara, CA) and subjected to direct sequencing at Eurofins Genomics K.K. (Tokyo, Japan). No indels or SNPs were observed in sequence chromatograms of females; some males where heterozygous due to detected double peaks and shifts of sequence reads. By sequencing from both sides, it was possible to obtain the genotypes of males and females (***Figure 3—figure supplement 2***).

### FISH analysis

In most lepidopteran species a conspicuous heterochromatic body is exclusively found in female polyploid nuclei. Since W derived-BAC as well as genomic probes have highlighted the W chromosomes and heterochromatin bodies in *B. mori* [45,46], there is no doubt that the bodies consist of the W chromosomes. The diagnosis however retains unreliable if a species of interest carries a W–autosomal translocation and/or partial deletion of the W [47,48]. Hiroki et al. [10] as well as Narita et al. [12] relied on the W-body diagnosis for C and CF females and concluded that they have WZ and ZZ sex chromosome constitutions, respectively. However, Kern et al. [13] has recently found that, on the basis of genomic qPCR designed to amplify Z-linked gene sequences (*Tpi* and *Ket*) relative to an autosomal gene (*EF-1α*), both CF and C females have only one Z chromosome while males have two Z chromosomes. This finding rejected the previous conclusion that the sex chromosome constitution of CF females is ZZ [10,12] but was inconclusive about whether CF females have a Z0 or W’Z system (with W’ as a modified W that has lost the feminization function and cannot be detected by the W-body assay). Hence we carried out more extensive chromosome analysis (other than just the W-body) to directly prove whether CF females carry the W or not.

In Lepidoptera, the W chromosome can be highlighted by FISH using probes prepared from whole genomic DNA of males or females. The capablity of FISH probes in detecting the W chromosome is due to the numerous repetitive short sequences occupying the W chromosome, which is then prone to be hybridized by random sequences. Genomic probes also paint repetitive regions scattered across other chromosomes, albeit at a lower density (autosomes and Z chromosome). Here we made mitotic and pachytene chromosome preparations from wing discs and gonads, respectively, in the last instar larvae of C and CF individuals of *E. mandarina* (see [49] for details). Genomic DNA was extracted from tet-treated C female larvae. Insect telomeric repeats were amplified by non-template PCR [50]. *Kettin* (*Ket*) gene fragments were amplified from adult cDNA synthesized by PrimeScript™ RT reagent Kit (TaKaRa, Otsu, Japan) and cloned by TOPO^®^ TA Cloning^®^ Kit (Thermo Fisher Scientific, Waltham, MA). We used 4 pairs of primers, Em_kettin_F1:5′–AGGTAATCCAACGCCAGTCG–3′ and Em_kettin_R1: 5′–TGCTTGCCCTAAGGCATTGT–3′, Em_kettin_F2: 5′–ACAATGCCTTAGGGCAAGCA–3′ and Em_kettin_R2: 5′–TGGGCAAAGCCTCTTCATGT–3′, Em_kettin_F3: 5′–AGATTCCGCACTACGCATGA–3′ and Em_kettin_R3: 5′–TAAATTGTGGTGGGACGGCA–3′, Em_kettin_F5: 5′–ACATGAAGAGGCTTTGCCCA–3’ and Em_kettin_R5: 5′–TCATGCGTAGTGCGGAATCT–3′, for PCR amplification with 94°C for 5 min followed by 35 cycles of 94°C for 30 s, 60°C for 30 s and 72°C for 3 min finalized by 72°C for 10 min. Probe labeling was done by using the Nick Translation Kit (Abbott Molecular, Des Plaines, IL). We selected Green-dUTP, Orange-dUTP (Abbott Molecular Inc.) and Cy5-dUTP (GE Healthcare Japan, Tokyo) fluorochromes for genomic DNA, *Ket* and insect telomeric repeat (TTAGG)*n* probes respectively. Hybridizations were carried out according to protocols described elsewhere [49]. Signal and chromosome images were captured with a DFC350FX CCD camera mounted on a DM 6000B microscope (Leica Microsystems Japan, Tokyo) and processed with Adobe Photoshop CS2. We applied green, red and yellow pseudocolors to signals from Green, Orange and Cy5 respectively.

### Quantitative polymerase chain reaction (qPCR)

Embryos of mated females were sampled 48 h after the oviposition and stored at –80°C until DNA extraction. Embryos were individually subjected to DNA extraction using DNeasy^®^ Blood & Tissue Kit (Qiagen, Tokyo, Japan). Real-time fluorescence detection quantitative PCR (qPCR) was performed using SYBR Green and a LightCycler^®^ 480 System (Roche Diagnostics K.K., Tokyo, Japan). Z-linked *Tpi* was amplified using TPI-F (5′–GGCCTCAAGGTCATTGCCTGT–3′) and TPI-R (5′–ACACGACCTCCTCGGTTTTACC–3′), Z-linked *Ket* was amplified using Ket-F (5′–TCAGTTAAGGCTATTAACGCTCTG–3′) and Ket-R (5′–ATACTACCTTTTGCGGTTACTGTC–3′), and autosomal *EF-1α* was amplified using EF-1F (5′–AAATCGGTGGTATCGGTACAGTGC–3′) and EF-1R (5′–ACAACAATGGTACCAGGCTTGAGG–3′) [13]. For each qPCR, a standard dilution series of PCR products (10^8^, 10^7^, 10^6^, 10^5^, 10^4^ and 10^3^ copies per microliter) was included in order to estimate the absolute copy numbers of the target sequence in the samples. To prepare standard samples, PCR products were gel-excised and purified by Wizard^®^ SV (Promega). Copy numbers of the standard samples were estimated by the concentration measured by a spectrophotometer, considering that the molecular weight of a nucleotide is 309 g/mol. For each qPCR, two replicates were performed that delivered similar results. All qPCRs were performed using a temperature profile of 40 cycles of 95°C for 5 s, 60°C for 10 s, and 72°C for 10 s. The qPCR data were analyzed by the Absolute Quantification analysis using the Second Derivative Maximum method implemented in the LightCycler^®^ 480 Instrument Operator Software Version 1.5 (Roche).

### RT-PCR

RNA was extracted from adult abdomens that were stored at -80°C using RNeasy^®^ Mini Kit (Qiagen, Tokyo, Japan). The cDNA synthesized by using Superscript™ III (Invitrogen) and Oligo(dT) was used as a template for RT-PCR. A partial sequence of *dsx* which contains alternative splicing sites was amplified using a primer pair, E520F (5′–GCAACGACCTCGACGAGGCTTCGCGGA–3′) and EhdsxR4 (5′–AGGGGCAGCCAGTGCGACGCGTACTCC–3′) and a temperature profile of 94°C for 2 min, 30 cycles of 94°C for 1 min, 57°C for 1 min and 72°C for 1 min 30 s, followed by 72°C for 7 min. The sequences of seven *dsx^F^* isoforms and a *dsx^M^* isoform were deposited in DDBJ/EMBL/Genbank (LC215389-LC215396).

## Acknowledgements

We thank Isao Kobayashi for help in collecting butterflies and Ranjit Kumar Sahoo for stimulating discussion. A part of this study is supported by NIAS technical support system at NARO.

## Additional information

### Funding

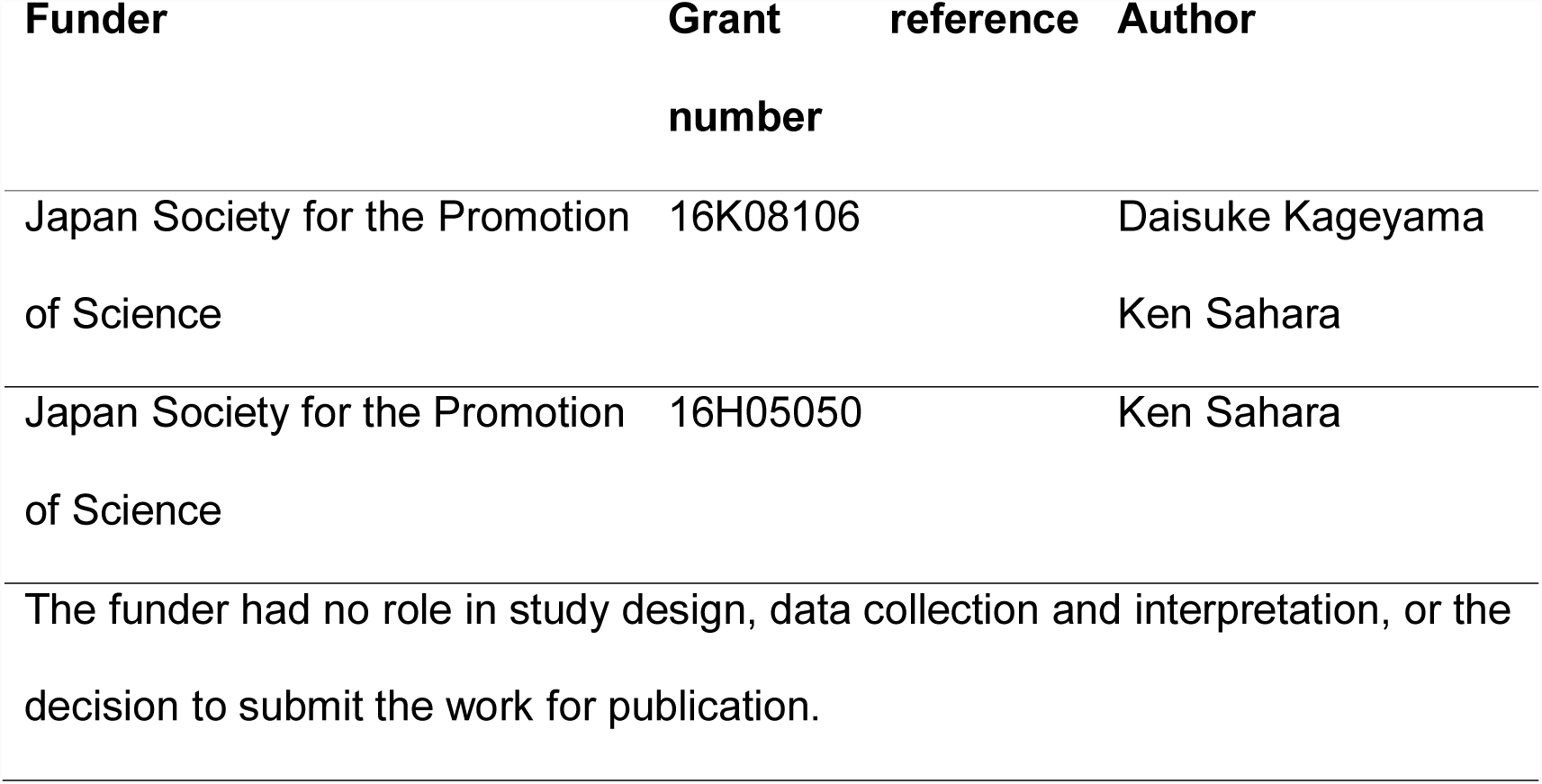

### Author contributions

DK, KS, designed the research; DK, MO, TS, AY, TK, SK, HK, YK, SN, MM, MR, KS, performed the research; DK, AJ, KS, analyzed the data; DK, MR, KS, wrote the paper with input from AY.

## Legends of figure supplements

**Figure 1**

**Figure supplement 1.**
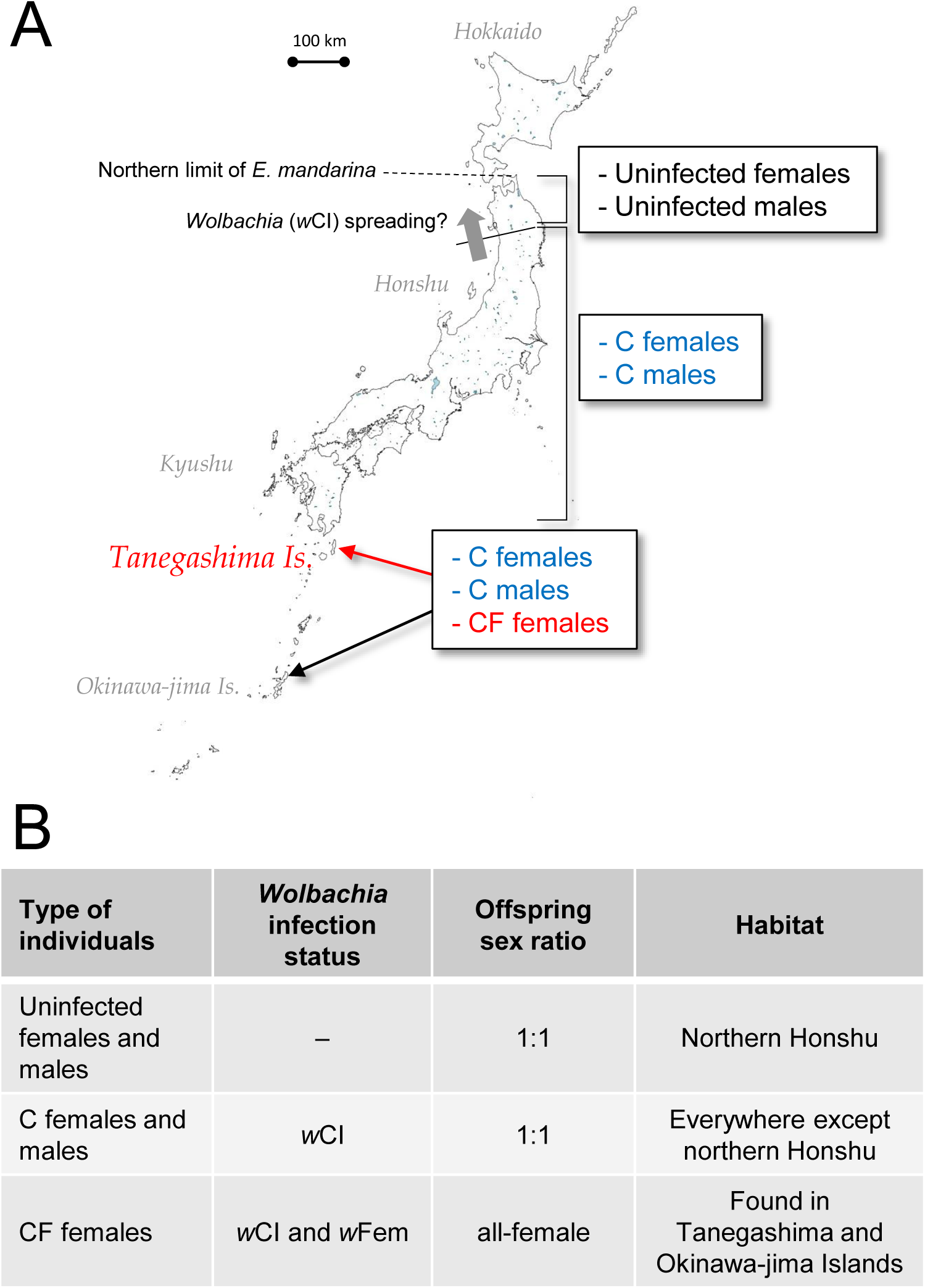
Habitat of *E. mandarina* in Japanese archipelago. (**A**) In this study, female adults of *E. mandarina* were collected on Tanegashima Island (map), located ca. 40 km from the southern tip of Kyushu, Japan. Within *E. mandarina*, the *Wolbachia* strain *w*CI is currently spreading northwards [8] together with the mitochondrial haplotypes introgressed from a sibling species (*E. hecabe*) by hybridization (hitchhiking effect; [9]). (**B**) On the basis of *Wolbachia* infection status, *E. mandarina* females can be categorized into three groups: uninfected females, C females (those singly infected with *w*CI), and CF females (those doubly infected with *w*CI and *w*Fem). These designations and their offspring sex ratio are summarized in the table. To date, in *E. mandarina*, CF females have only been found on Okinawa-jima Island [10,11] and Tanegashima Island [12,39].

**Figure 2**

**Figure supplement 1.**
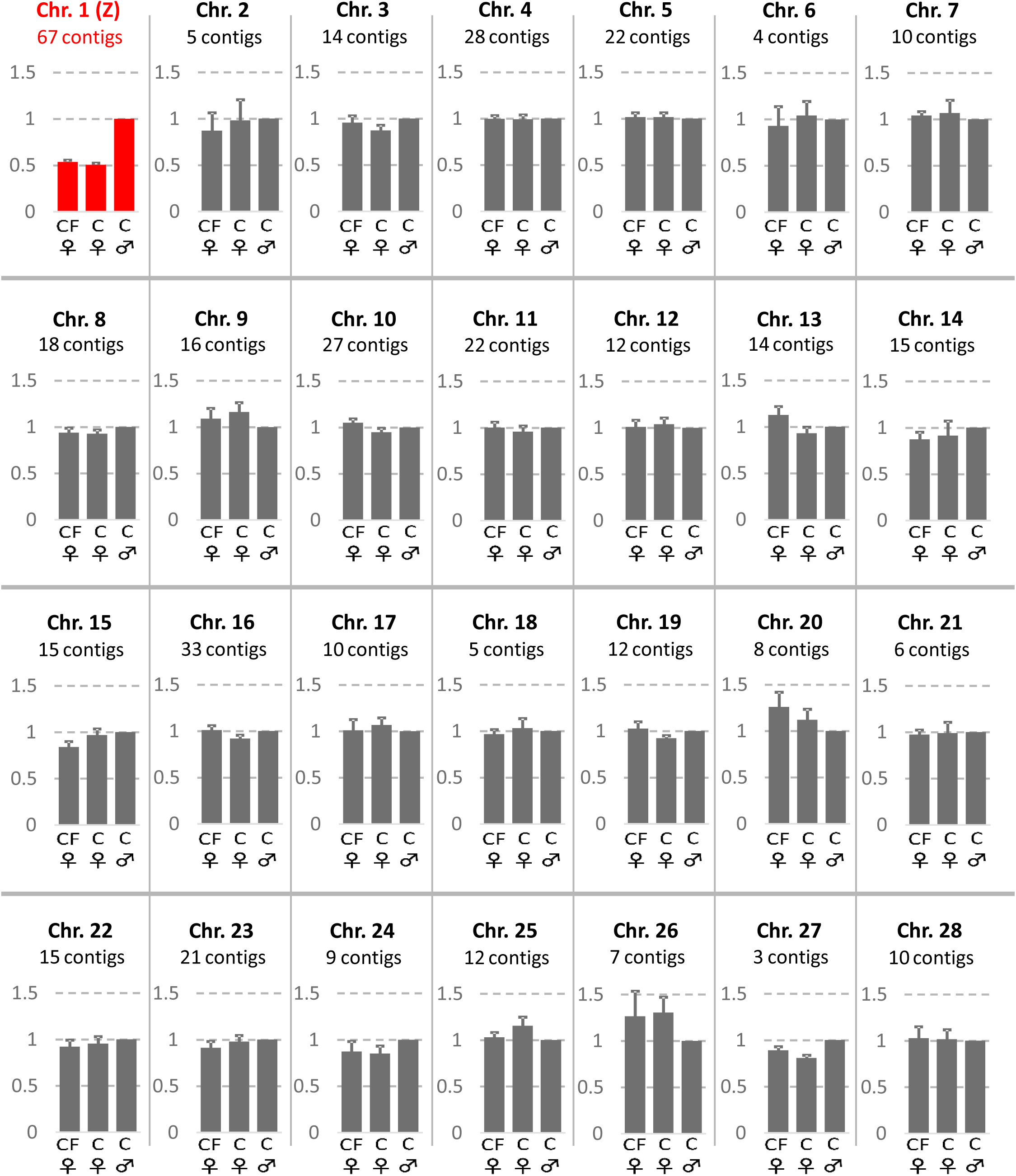
Relative normalized sequence read counts for 440 contigs of *E. mandarina* that matched to *B. mori* loci on 28 chromosomes. Means and standard errors are shown for CF females and C females while those of C males were set to 1.

**Figure 3**

**Figure supplement 1.**
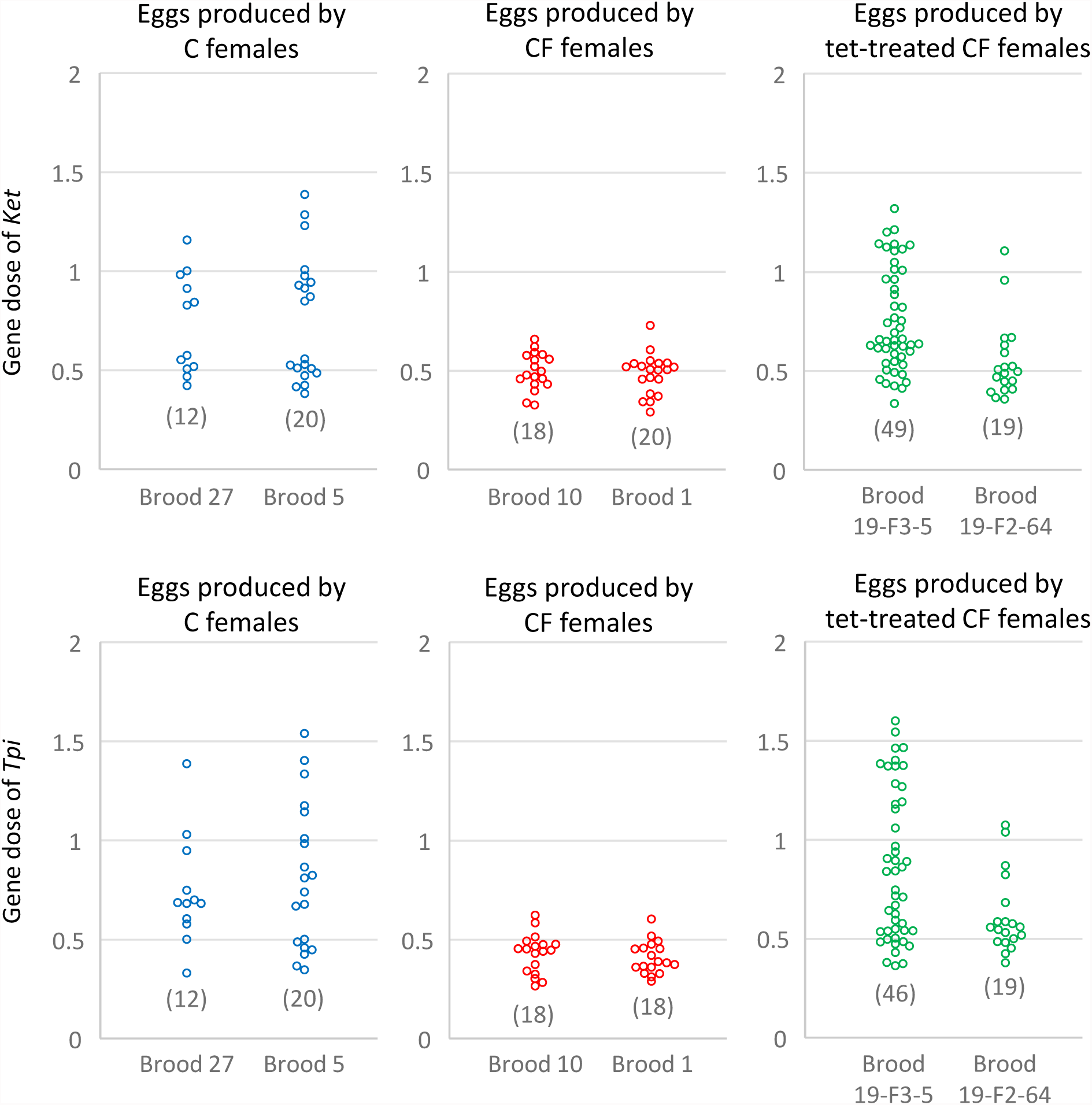
Estimate of Z-linked gene dose of *E. mandarina*. Estimate of the gene dose of *Ket* (top) and *Tpi* (bottom), relative gene copies per *EF-1α*, by genomic qPCR in each of the fertilized eggs laid by C females, CF females and tet-treated CF females. Each circle represents an egg. Each of the codes along the x-axes indicate the brood produced by a single mother.

**Figure supplement 2.**
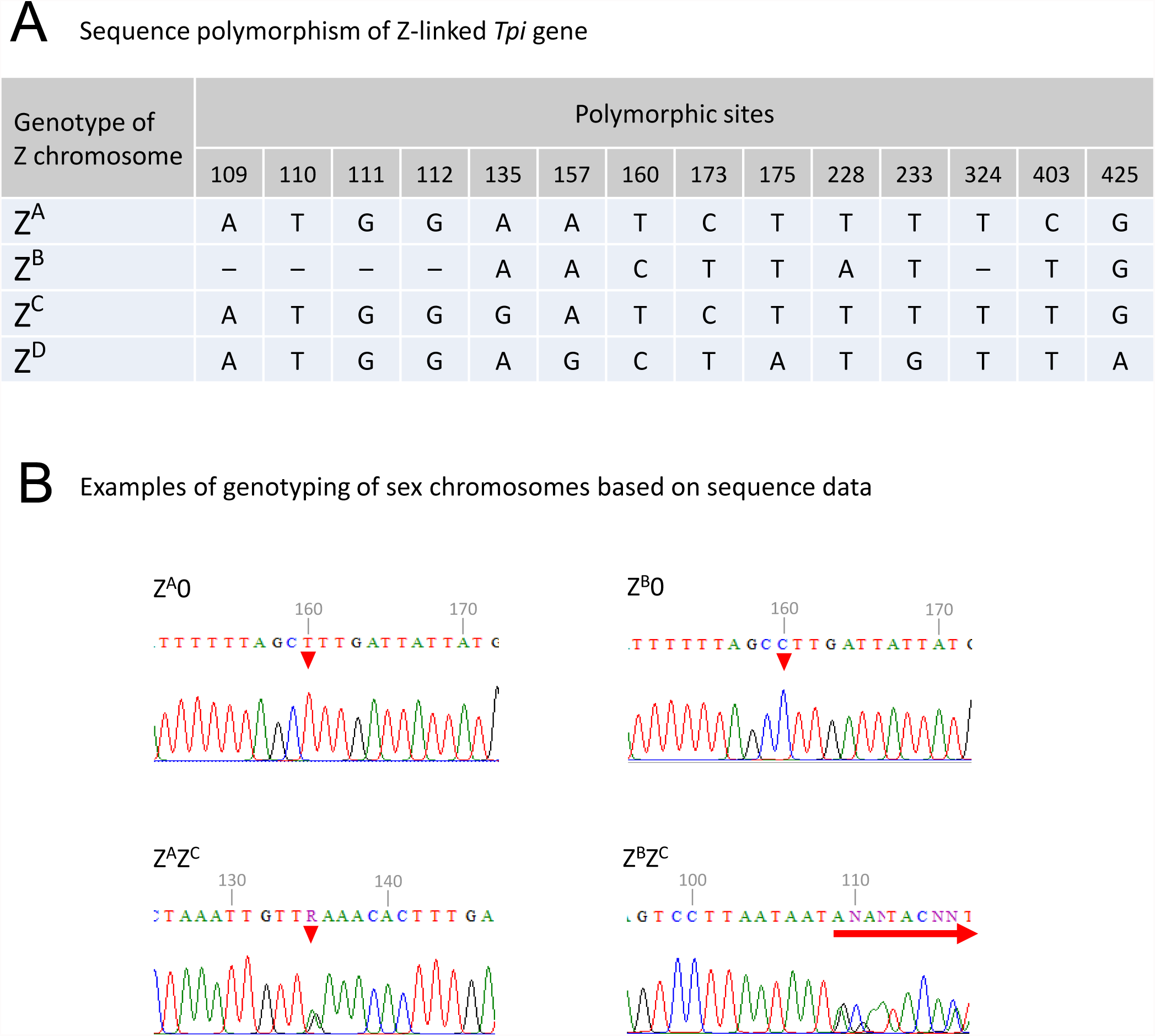
Genotyping of Z chromosome based on nucleotide polymorphism of *Tpi*. (**A**) Sequence polymorphism of *Tpi*. In our experiment, Z chromosomes were categorized into four (Z^A^, Z^B^, Z^C^ and Z^D^) on the basis of *Tpi* sequence. An en dash represents a gap. (**B**) Examples of genotyping based on *Tpi* sequence data. Red triangles represent polymorphic sites. When Z^B^ was paired to Z^A^, Z^C^ or Z^D^, sequence gaps resulted in ambiguity from the position 109 (shown with a red arrow).

**Figure 4**

**Figure supplement 1.**
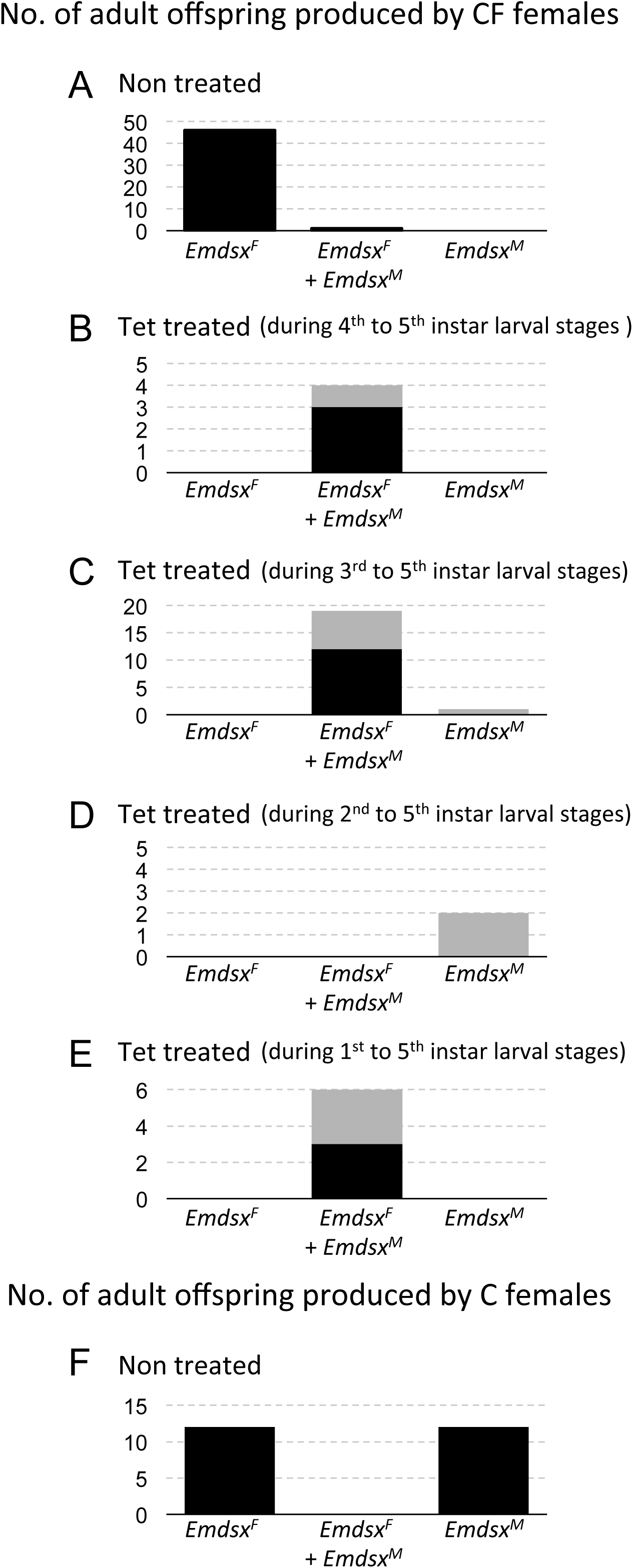
Detection of *Emdsx* in adults that were tet-treated during various larval stages. The numbers of adults that failed to emerge from their pupal cases are shown with gray.

**Figure supplement 2.**
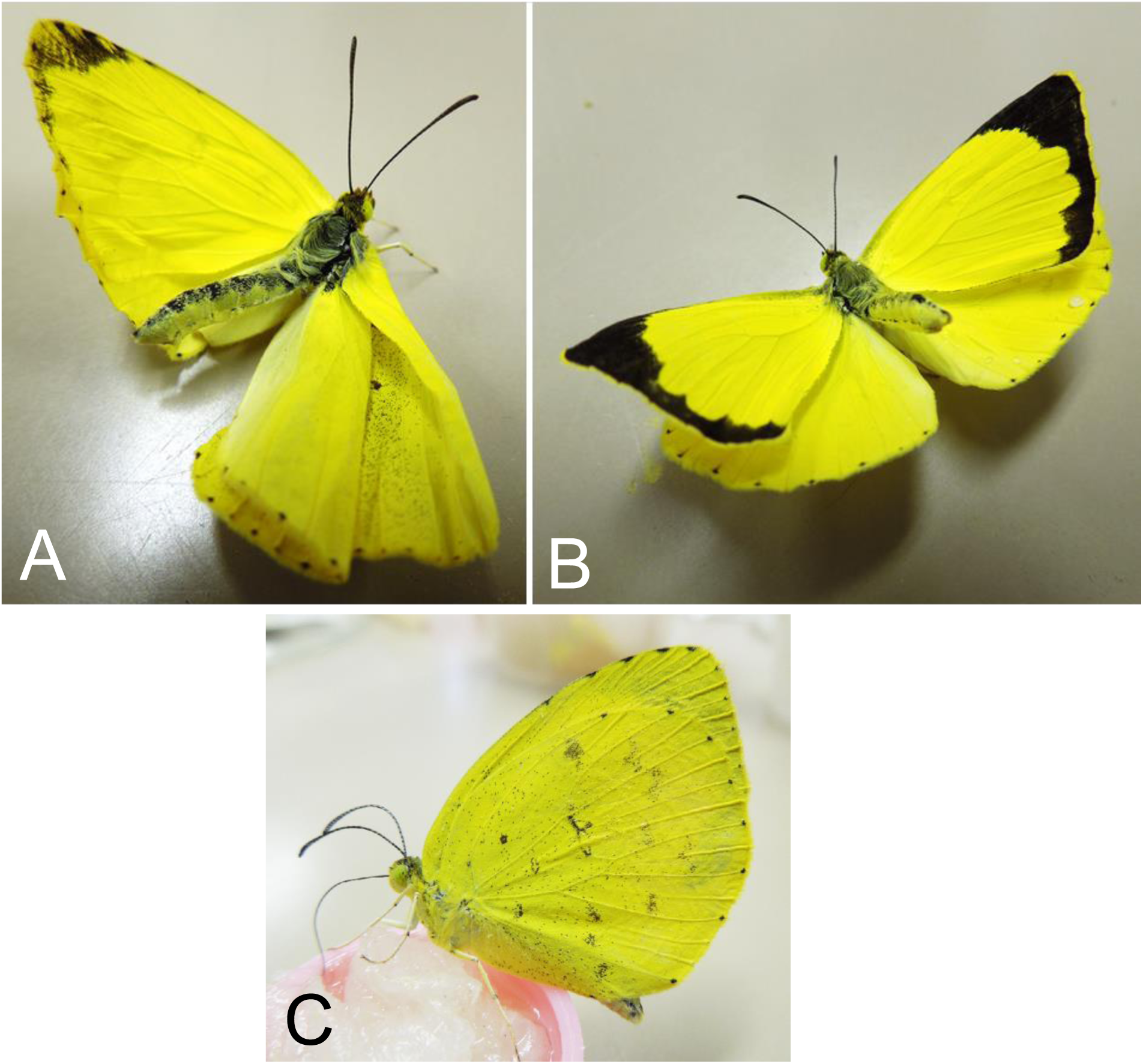
(**A**-**B**) Intersexual adults generated by feeding the CF larvae with tet-containing diet. Their wings are often curled or crumpled. Most of them are trembling and cannot stand still. (**C**) Normal females. Their wings are neatly closed.

**Figure supplement 3.**
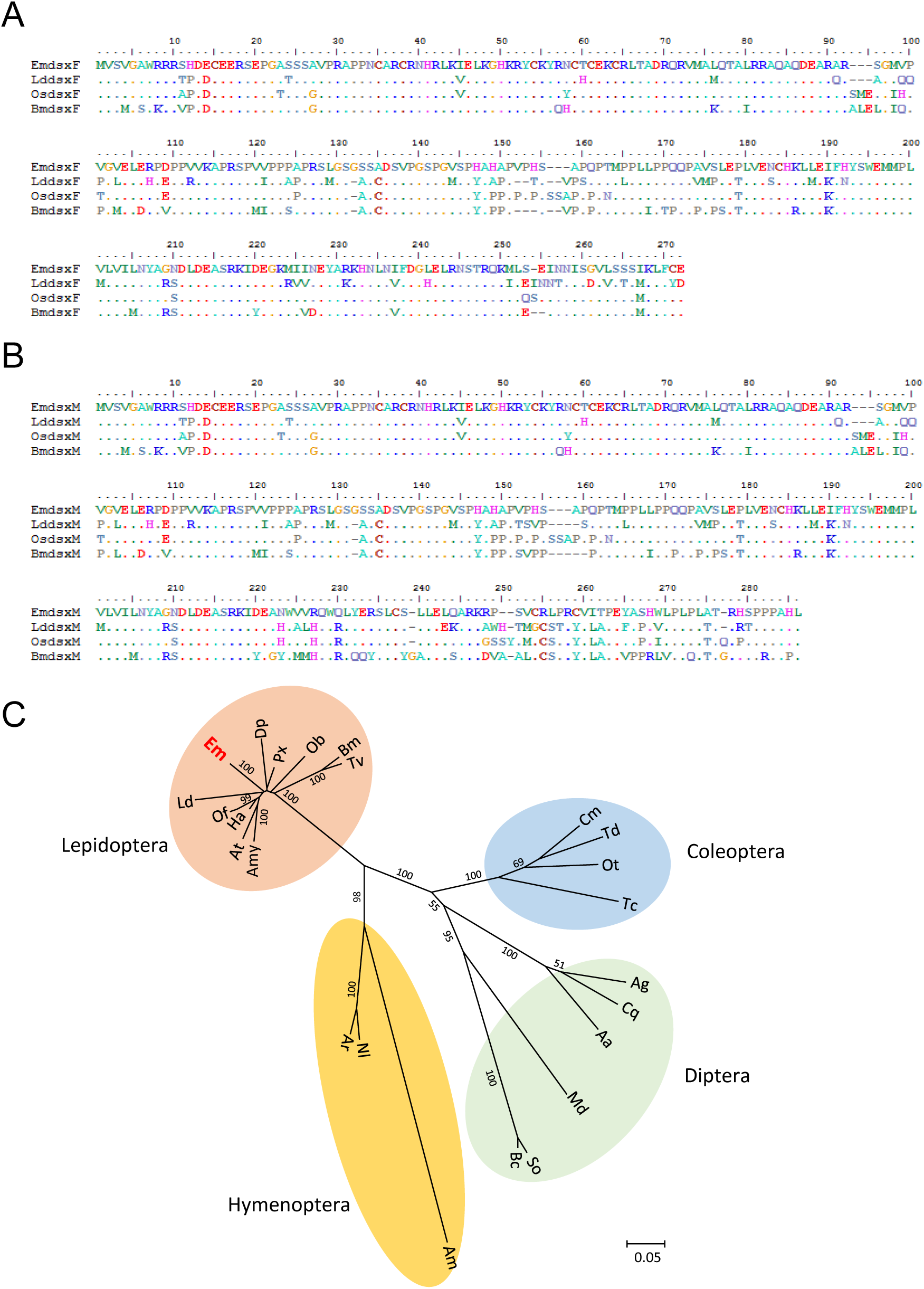
(**A**) Amino acid sequences of female splice forms of *dsx* genes derived from *Eurema mandarina* (*Emdsx^F^*: LC215389) and other lepidopteran species, *Lymantria dispar* (*Lddsx^F^*: BAN82533), *Ostrinia scapulalis* (*Osdsx^F^*: BAJ25851) and *Bombyx mori* (*Bmdsx^F^*: NP_001036871). (**B**) Amino acid sequences of male splice forms of *dsx* genes derived from *E. mandarina* (*Emdsx^M^*: LC215396), *L. dispar* (*Lddsx^M^*: BAN82532), *O. scapulalis* (*Osdsx^M^*: BAJ25850), and *B. mori* (*Bmdsx^M^*: AHF81625). (**C**) Unrooted NJ tree of the *dsx* gene based on amino acid sequences. Em: *E. mandarina* (LC215389), Dp: *Danaus plexippus* (EHJ78146), Px: *Papilio xuthus* (XP_013171086), Ob: *Operophtera brumata* (KOB69684), Bm: *B. mori* (NP_001036871), Tv: *Trilocha varians* (BAS02078), Amy: *Antheraea mylitta* (ADL40853), At: *Amyelois transitella* (XP_013184257), Ha: *Helicoverpa armigera* (AHF81652), Of: *Ostrinia furnacalis* (AHF81640), Ld: *L. dispar* (BAN82533), Am: *Apis mellifera* (ABV55180), Nl: *Neodiprion lecontei* (XP_015517992), Ar: *Athalia rosae* (XP_012262273), Cm: *Cyclommatus metallifer* (BAO23810), Td: *Trypoxylus dichotomus* (BAM93344), Ot: *Onthophagus taurus* (AEX92939), Tc: *Tribolium castaneum* (AFQ62107), Ag: *Anopheles gambiae* (XP_309601), Cq: *Culex quinquefasciatus* (AJB28478), Aa: *Aedes aegypti* (ABD96571), Md: *Mayetiola destructor* (AGW99160), So: *Sciara ocellaris* (CDN30082), Bc: *Bradysia coprophila* (CDN30080).

